# Siglec-F Protects Against Elastase-induced Lung Inflammation and Emphysema in Mice

**DOI:** 10.1101/2025.09.16.674299

**Authors:** Qihua Ye, Naoko Hara, Yan Hu, Marika Orlov, Bradford J. Smith, Tonya M Brunetti, Rachel Z Blumhagen, Zhenghui Liu, Bifeng Gao, Melanie Königshoff, William J. Janssen, Christopher M. Evans

## Abstract

Respiratory surfaces are exposed daily to billions of inhaled particles that could cause injury to alveolar tissues where gas exchange occurs. Accordingly, robust defense is essential but must involve minimal physiologic disruption. Airspace macrophages (AMs) protect lung surfaces while also maintaining tissue integrity through non-inflammatory homeostatic responses in health. However, in chronic obstructive pulmonary disease (COPD), AMs contribute to emphysematous alveolar destruction through mechanisms that are poorly understood. We hypothesized that damaging effects of AMs in emphysema result from loss of mechanisms that normally restrain homeostatic AMs. We found that Siglec-F, a common marker of AMs, exerts suppressive effects on tissue resident AMs (RAMs), thereby supporting alveolar integrity in mouse lungs during health and in a model of elastase-induced alveolar destruction. Siglec-F-deficient mice exhibited decreased alveolar numbers at baseline, and they had worsened alveolar damage after elastase challenge that was unexpectedly RAM-mediated. Transcriptomic profiling revealed dysregulation of key pathways involved in tissue remodeling and repair, including extracellular matrix degradation, TGF-β signaling, and phagocytosis. These findings uncover previously unrecognized roles for RAMs and Siglec-F in preserving alveolar integrity. Our findings implicate RAMs and Siglecs as potential therapeutic targets for preserving alveolar integrity in health and for limiting alveolar damage in COPD.

## Introduction

Every day, the lungs are exposed to myriad particles and pro-inflammatory materials that could potentially damage delicate alveolar gas exchange surfaces. Healthy lungs thus rely on protective epithelial barriers and airspace macrophages (AMs) as critical lines of host defense (1). During health, AMs serve as sentinels that minimize inflammation while preserving tissue integrity (2). Following injury, AMs help restore homeostasis and promote lung repair. However, AMs can also worsen lung injury in pathologic settings such as chronic obstructive pulmonary disease (COPD) (3).

A major manifestation of COPD is emphysema, characterized by airspace enlargement and impaired gas exchange due to loss of alveoli (4). Key drivers of this pathology are alveolar epithelial cell damage and destruction of the underlying extracellular matrix by proteases (5). AMs are well recognized as major sources of proteases that contribute to alveolar destruction in COPD (3). Nonetheless, mechanisms regulating homeostatic versus detrimental functions of AMs remain poorly understood. The current conceptualization of these differences centers on AM ontogeny.

During health, lungs are occupied by embryonic-derived resident AMs (RAMs) that perform homeostatic functions. After injury, RAMs are joined by recruited AMs (RecAMs) that mature from monocytes that migrate to sites of damage. These two AM subsets are typically considered functionally distinct. For example, RecAMs augment tissue damage during early phases of lipopolysaccharide (LPS) or bleomycin-induced lung injury, but RecAMs can contribute to repair at later time points (6–9). By contrast, RAM programming in these injury settings remains relatively static, and this may reflect continued restraint that was placed on RAMs during health to optimize homeostatic defense and alveolar function (primarily gas exchange).

To achieve homeostatic quiescence, RAMs express molecules that limit inflammatory signaling. <u>S</u>ialic acid-binding immuno<u>g</u>lobulin-like <u>lec</u>tins (Siglecs) are surface proteins on leukocytes that comprise numerous inhibitory immunoreceptors (10). In mice, Siglec-F is highly expressed by AMs and is thought to exert inflammosuppressive effects that promote homeostatic functions in health and facilitate repair after injury (6). Recently, we demonstrated that bleomycin-induced lung fibrosis in mice was alleviated by Siglec-F deficiency (11). However, underlying mechanisms through which Siglec-F mediates AM function are still only partially understood.

To further elucidate the role of Siglec-F in AM homeostasis and its potential involvement in COPD pathogenesis, we investigated the effects of Siglec-F deficiency in health and in a murine model of elastase-induced emphysema. We found that in the absence of Siglec-F, RAMs (but not RecAMs) accumulated, and RAMs adopted a detrimental phenotype that exacerbated alveolar destruction. This reveals a previously unknown capacity of RAMs to contribute to alveolar damage. Although RAMs are canonically viewed as pro-repair and homeostatic, our findings indicate that their programming is Siglec-F-dependent. Without Siglec-F, the lungs demonstrated mild alveolar damage in health, and worsened emphysema after elastase challenge that was directly linked to transcriptional reprogramming of RAMs toward a more tissue-destructive state. These results demonstrate that Siglec-F is not merely a marker. It is also a critical regulator of AM function whose its deficiency promotes COPD-like pathology in mice.

## Methods

Detailed methods are provided in an online supplement.

### Sex as a biological variable

Our study examined male and female animals, and similar findings are reported for both sexes.

### Animal model

C57BL/6J congenic *Siglecf^+/+^* and *Siglecf*^−/−^ mice were bred and housed at the University of Colorado and National Jewish Health. Mice were used starting at 12 weeks of age. One dose of oropharyngeal (o.p.) porcine pancreatic elastase (PPE) was administered to mice at a dose of 20 U/kg. Endpoints were assessed on days 0, 1, 3, 7, and 21.

### Lung physiology

Lung function was measured using a flexiVent (Scireq). Mice were anesthetized with urethane (2.0 g/kg, i.p.), tracheostomized, and ventilated (150 breaths/min, 10 ml/kg, 3 cmH_2_O positive end expiratory pressure). Oscillatory mechanics and pressure-volume (P-V) relationships were assessed as described previously (12).

### Lung morphometry

Stereology was performed in accordance with ATS/ERS standards (13). Briefly, lungs were inflated and fixed by intratracheal instillation of 1% low-melting agarose and 4% paraformaldehyde at 25 cmH₂O pressure. Fixed lungs were embedded in 3% agar with randomized orientations and sectioned into 2 mm slabs. Slabs were randomly selected for paraffin embedding, and 5 μm H&E-stained sections were prepared for stereological analysis. Total alveolar number and chord length were quantified to assess alveolar destruction and airspace enlargement, respectively.

### AM isolation, quantification, and sorting

Right lungs were lavaged after clamping the left lungs at the mainstem bronchus. The right lung was lavaged by instilling and removing 0.5 ml PBS containing 0.5 mM EDTA three times (1.5 ml total). Lung lavage was used for hemocytometer counts, Giemsa stain differentials, flow cytometry, and FACS.

### AM flow cytometry and sorting

Erythrocytes from lavage were lysed with RBC lysis buffer (eBioscience) for 10 minutes at room temperature. After washing, the cells were labeled with anti-CD16/32, CD45, CD64, CD88, Ly6G, CD11b, CD11c, and Siglec-F antibodies (all from BioLegend) at 4 °C for 45 min. The cells were centrifuged at 300 x g for 8 min, washed in FACS buffer, and re-suspended in 500 μl FACS buffer (BioLegend). Before flow cytometry or sorting, one drop of NucBlue (Invitrogen) or DAPI (BioLegend) was added to exclude dead cells. Neutrophils were identified as CD45^+^Ly6G^+^, AMs were gated as CD45^+^Ly6G^-^CD64^+^CD88^+^ cells, and recruited and resident AMs were separated based on CD11b and CD11c expression as previously reported (6, 14). FACS was performed on a Sony Biotechnology SY3200 sorter. Flow cytometry was performed on a BD LSR Fortessa analyzer.

### Bulk RNA-seq

RNA was extracted from sorted cells using a microRNA kit (Qiagen). Integrity and quantity were assessed via NanoDrop and TapeStation 4200. Libraries were prepared from 100 ng RNA using MGIEasy RNA Library Prep Set (3.1) and sequenced as 100 bp paired-end reads (∼ 80M reads/sample) on the DNBSeq-7000 (MGI). Reads were trimmed with cutadapt (4.2), aligned with STAR (2.7.10b), and strandedness was evaluated using Picardtools (2.27.5). Genes with >10 raw counts in ≥10 of 27 samples were retained. Normalization and differential expression were performed using DESeq2 (1.38.2) in R (4.2.2), and gene set enrichment analysis (GSEA) was done using fgsea (1.24.0).

### Statistics

Statistical analyses for non-RNA seq data were performed using GraphPad Prism (10.2.2). Two-sample comparisons were performed using unpaired two-tailed t-tests or Mann-Whitney U-tests where appropriate. For multiple comparisons, ANOVA with a Dunnett post-hoc correction or a Kruskal-Wallis test with Dunn’s post-hoc correction were used. For RNA-seq data, differentially expressed genes (DEGs) were identified using a threshold of adjusted p-value of <0.05 and log_2_FC of > 1 or < -1. Significant pathways identified through GSEA were defined by an adjusted p-value < 0.05 and normalized enrichment scores (NES) > 1.5 or < -1.5.

### Study approval

This study was approved by and performed in accordance with IACUC protocols at University of Colorado Anschutz Medical Campus and National Jewish Health.

### Data availability

Raw sequence files and processed files are available on the Gene Expression Omnibus (accession no. GSE307353). Codes for main analyses is available at https://github.com/QIHUA-YE. Values for all data points in graphs are reported in the Supporting Data Values file.

## Results

### Acute lung injury caused by porcine pancreatic elastase (PPE) leads to progressive emphysema in mice

To develop a model by which tissue protective effects of Siglec-F on AMs could be evaluated, we treated wild-type mice with intratracheal porcine pancreatic elastase (PPE). Consistent with previous studies (15–18), a single dose of PPE induced emphysema, as confirmed by histologic quantification of mean chord length measurements (Figures 1A, B). Airspace enlargement was apparent 3 days post-PPE and progressively worsened over a 21-day time course.

**Figure 1.**
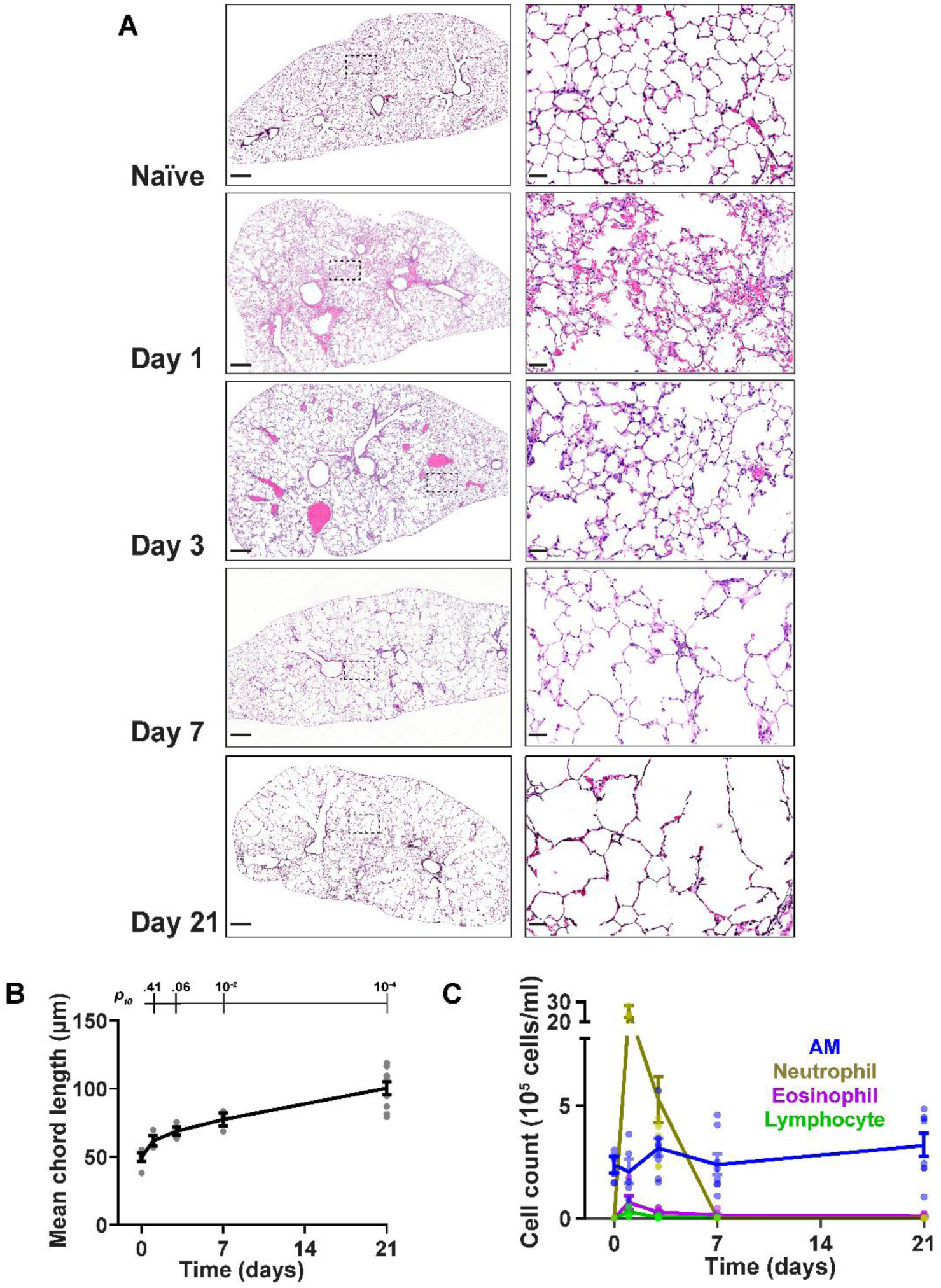
A single dose of porcine pancreatic elastase (PPE) induces lung inflammation and emphysema-like injury. **(A)** Representative lung histology images across time points. Scale bars: 500 μm (left) and 50 μm (right). The dashed box on the left indicates the region at a higher magnification on the right. **(B)** Mean chord length measurements across time points (n = 3-9 per time point). Statistical analysis was performed using one-way ANOVA with Dunnett’s multiple comparisons test; adjusted p-values for each time point relative to day 0 (*p_t0_*) are shown above the time curve. **(C)** Differential cell counts in lung lavage. (n = 4-10 per time point). Data are presented as mean ± SEM.

To assess inflammation, we quantified leukocytes in lung lavage. PPE induced robust and transient neutrophilia that peaked at day 1 and resolved by day 7 (Figure 1C, Table E1). Small numbers of eosinophils and lymphocytes were also detectable at early time points but resolved by day 7. By contrast, AMs were present in substantial numbers at all time points and predominated from day 7 onward as alveolar enlargement progressed. Accordingly, we sought to determine how PPE-induced emphysema may be causatively related to AMs.

### Resident AMs predominate in the airspace during PPE-induced lung inflammation and injury

To characterize the kinetics of AM subsets during PPE-induced inflammation, we performed flow cytometry on cells from lung lavage (Figure 2A). Leukocytes were identified with CD45, and neutrophils were distinguished as Ly6G⁺. AMs were identified as Ly6G^-^ CD64^+^ CD88^+^ cells and defined as resident AMs (RAMs) based on CD11c expression or recruited AMs (RecAMs) based on CD11b expression (6). Acute lung injury resulted in migration of RecAMs into the lungs on days 1, 3, and 7 after PPE challenge. However, throughout the time course, RAMs were more abundant than RecAMs, with significant differences observed at most time points (Figure 2B).

**Figure 2.**
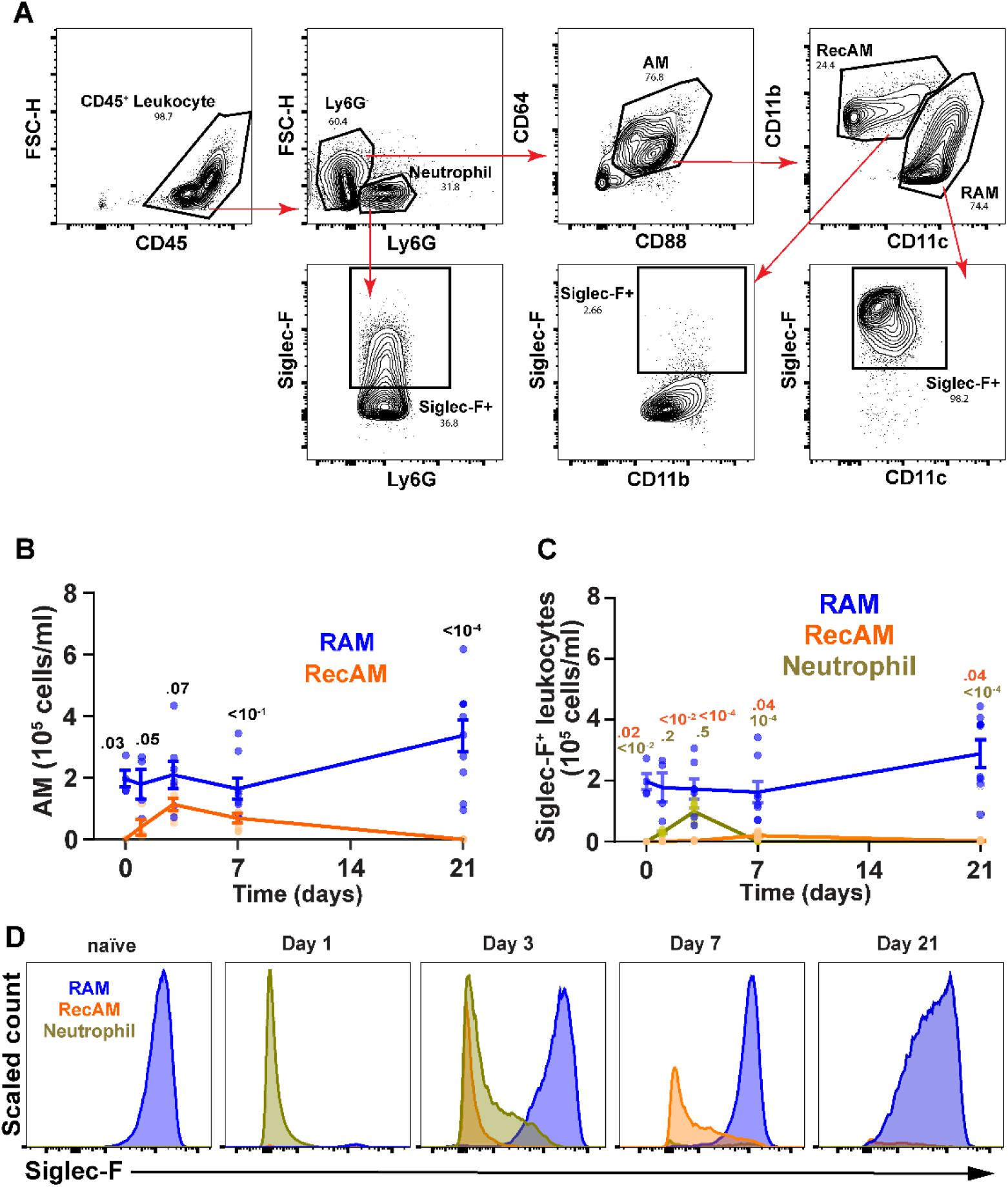
CD11c⁺ Siglec-F⁺ RAMs predominate during PPE-induced lung inflammation. **(A)** Representative gating strategy for identifying immune cells in lung lavage (day 3 shown). **(B)** Quantification of alveolar macrophage (AM) subsets over time. Mann-Whitney test was performed at each time point to compare RAM and RecAM populations; adjusted p-values for these comparisons are shown. **(C)** Quantification of Siglec-F⁺ cells in lung lavage. Kruskal-Wallis test with Dunn’s multiple comparisons was performed at each time point; adjusted p-values for RecAM and neutrophils relative to RAM are indicated. (n = 4-10 per time point). Data are presented as mean ± SEM. **(D)** Siglec-F expression across major cell populations. Cell count was scaled to show the maximum at each time point.

Quantification of Siglec-F⁺ cells in lung lavage revealed that RAMs consistently represented the major Siglec-F⁺ immune cell population (Figure 2C). In contrast, recruited AMs (RecAMs) expressed minimal Siglec-F at all time points. A subset of neutrophils (∼30%) transiently expressed Siglec-F on day 3, but expression levels were lower than those observed in RAMs (Figures 2A, D). To ensure that PPE administration did not disrupt Siglec-F on AMs, we tested whether Siglec-F detection is affected by PPE. AMs were treated with PPE *in vivo* and *in vitro* for 1 h. Siglec-F expression on AMs remained intact (Figure E1). Thus, Siglec-F is stably expressed on RAMs at a high level that is not substantially impacted by PPE’s proteolytic activities. Collectively, these data demonstrate that the inflammatory response to PPE challenge is typified by a Siglec-F-expressing RAM population. We next sought to determine the functional significance of Siglec-F expression.

### Absence of Siglec-F worsens emphysema

To investigate the role of Siglec-F in PPE-induced emphysema, we compared responses to PPE challenge in Siglec-F knockout (*Siglecf^-/-^*) mice and their wild-type (*Siglecf^+/+^*) littermates. Lung function was measured at baseline and on day 21 post-PPE. We observed significantly increased static compliance (Cst), inspiratory capacity, and total lung volume, as well as significantly decreased respiratory system and pulmonary system elastance in *Siglecf ^-/-^* mice on day 21 compared to WT mice (Figures 3A-C, E2A-B), consistent with more severe emphysema. Pressure-volume loops confirmed increased inspiratory capacity in *Siglecf^-/-^* mice, illustrated by an upward shift of the loop compared to *Siglecf^+/+^* (Figure E2C). There were no differences in airway and tissue resistance (Figure E2).

**Figure 3.**
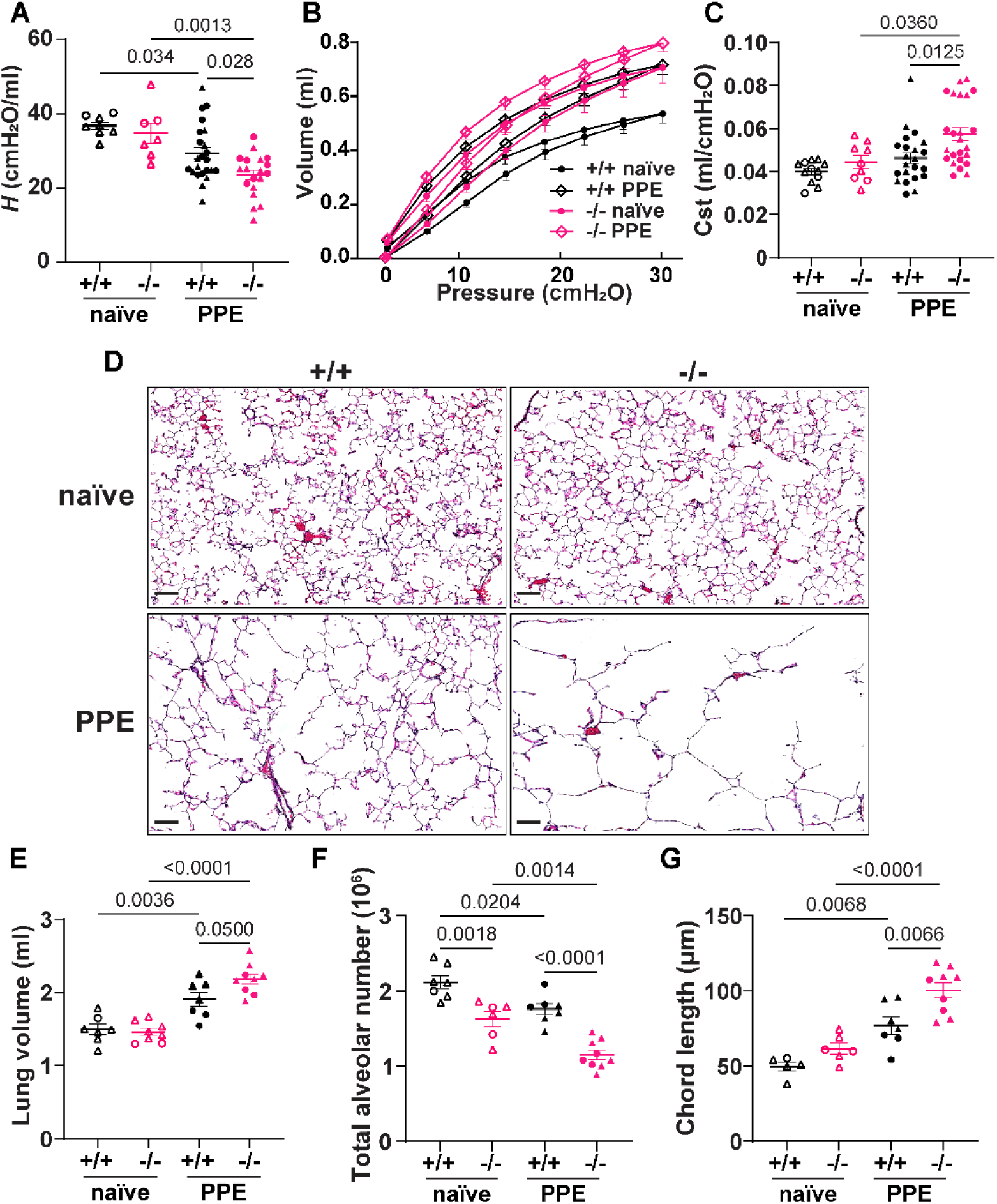
Siglec-F deficiency exacerbates PPE-induced COPD-like injury. **(A)** Lung tissue elastance (*H*), **(B)** Pressure-Volume loop, **(C)** static compliance (Cst)**, (D)** Representative lung histology images, Scale bars: 100 μm. **(E)** total lung volume determined by Archimedes principle, **(F)** Chord length, and **(G)** total alveolar number quantification in *Siglecf^+/+^*and *Siglecf^-/-^* mice at baseline and day 21 post-PPE. Triangles denote female mice; circles denote male mice. Data are shown as mean ± SEM. One-way ANOVA with Dunnett’s post hoc test was performed.

To evaluate the effects of *Siglecf* deletion on lung structure, unbiased stereology was used to quantify alveolar number and chord length on lungs from *Siglecf^-/-^* mice and *Siglecf^+/+^* littermates at baseline and 21 days after PPE. In line with the visual appearance of histologic sections (Figure 3C), alveolar numbers were reduced and chord length was increased in *Siglecf^-/-^* compared to *Siglecf^+/+^* mice after PPE-induced lung injury (Figures 3E, F).

Although there were no differences in compliance, inspiratory capacity, total lung volume, elastance, or chord length between *Siglecf^-/-^*and *Siglecf^+/+^* mice at baseline (Figures 3A-C, F, E2A-B), we observed significant decreases in alveolar numbers in *Siglecf^-/-^* mice (Figure 3E). To investigate whether these baseline differences might be due to defects in alveologenesis, we quantified alveolar numbers in post-natal day 14 *Siglecf^+/+^* and *Siglecf^-/-^*mice. There were no significant differences at this key time point of lung development (Figure E4), suggesting that the difference observed in adults reflects a homeostatic role for Siglec-F that becomes even more apparent after injury.

### Siglec-F deficiency leads to the accumulation of AMs, primarily RAMs

To specifically address how Siglec-F influences immune cell dynamics following PPE-induced injury, we compared leukocyte populations in *Siglecf^-/-^*and *Siglecf^+/+^* mice. At baseline, we observed a significant increase in total AMs, which were all CD11c^+^ RAMs, in *Siglecf^-/-^*mice compared to *Siglecf^+/+^* mice. There were no differences in other cell types (Figure 4A-B, Tables E2-3). On day 7 post-PPE, we observed a significant increase in total AMs (Figure 4A), largely driven by RAMs (Figure 4B) in *Siglecf^-/-^*. RecAMs did not show a significant change (Figure 4C, Tables E2-3).

**Figure 4.**
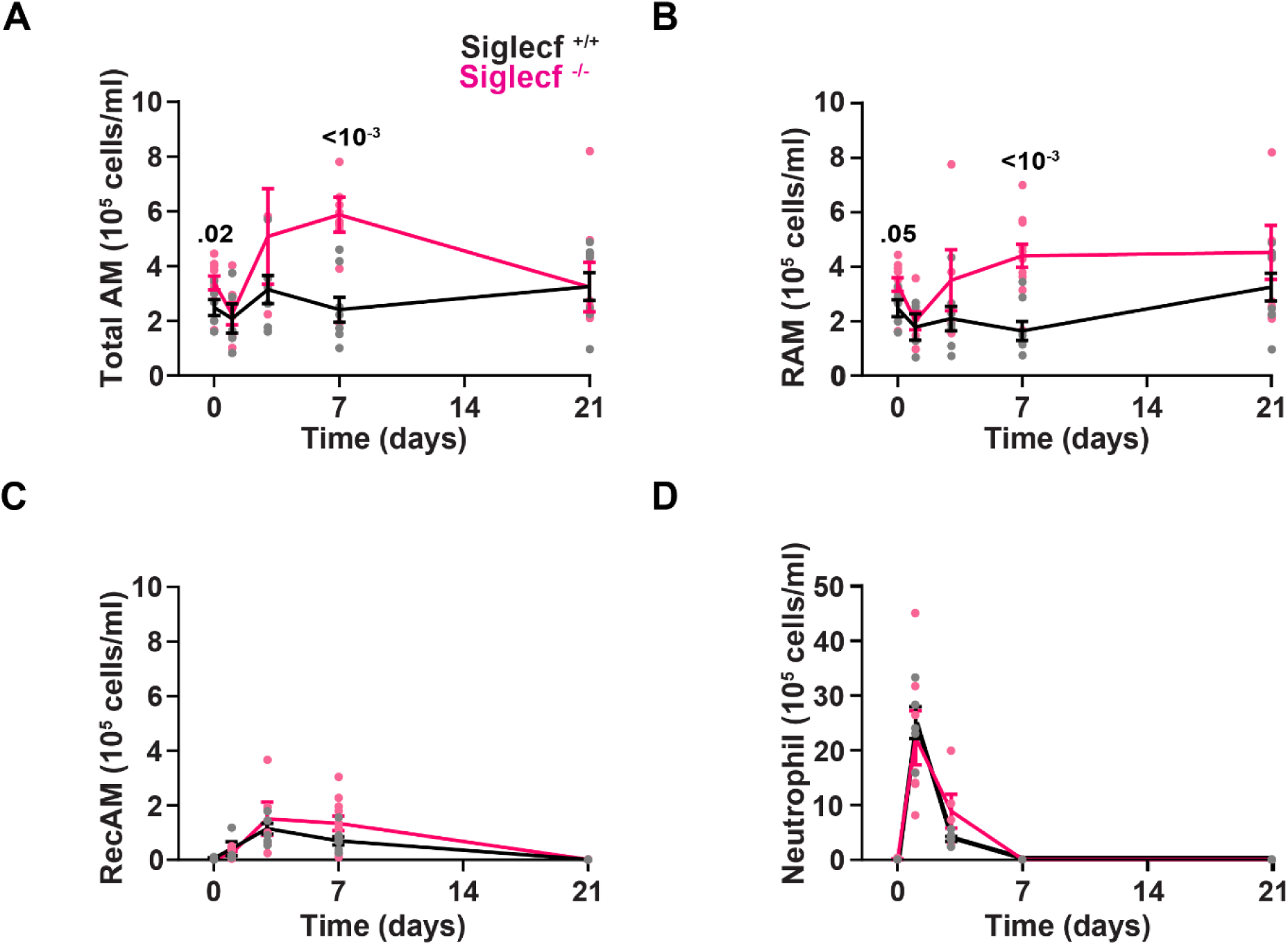
Siglec-F deficiency leads to the accumulation of AMs, primarily CD11c+ RAMs, on day 7. Quantification of **(A)** total AM, **(B)** RAM, **(C)** RecAM, and **(D)** neutrophil in lung lavage from *Siglecf^+/+^* and *Siglecf^-/-^* mice (n = 4-10 per time point). Data are shown as mean ± SEM. Statistical analyses between *Siglecf^+/+^* and *Siglecf^-/-^* mice at each time point were performed using Mann-Whitney test. Significant adjusted p-values are indicated.

Total neutrophil numbers were not significantly different between genotypes, despite the moderate Siglec-F expression in a small subset of neutrophils previously observed in *Siglecf^+/+^*mice on day 3 (Figures 4D, 2A-B, Tables E2-3). Eosinophils also remained low in number and showed no significant differences between genotypes at any time point (Figure E5, Tables E2-3).

Collectively, these results indicate that loss of Siglec-F promotes the expansion of RAMs in the alveolar space, underscoring its role in modulating RAM dynamics and the inflammatory milieu following PPE-induced injury.

### Siglec-F deficiency leads to transcriptional reprogramming of RAM effector functions

To investigate whether effector functions of RAMs were perturbed in the absence of Siglec-F, we performed bulk RNA-seq on sorted CD11c⁺ RAMs isolated from *Siglecf^-/-^*and *Siglecf^+/+^* mice on days 0, 3, 7, and 21. Gene set enrichment analysis (GSEA) revealed pathways perturbed by the absence of Siglec-F in RAMs (Figure 5).

**Figure 5.**
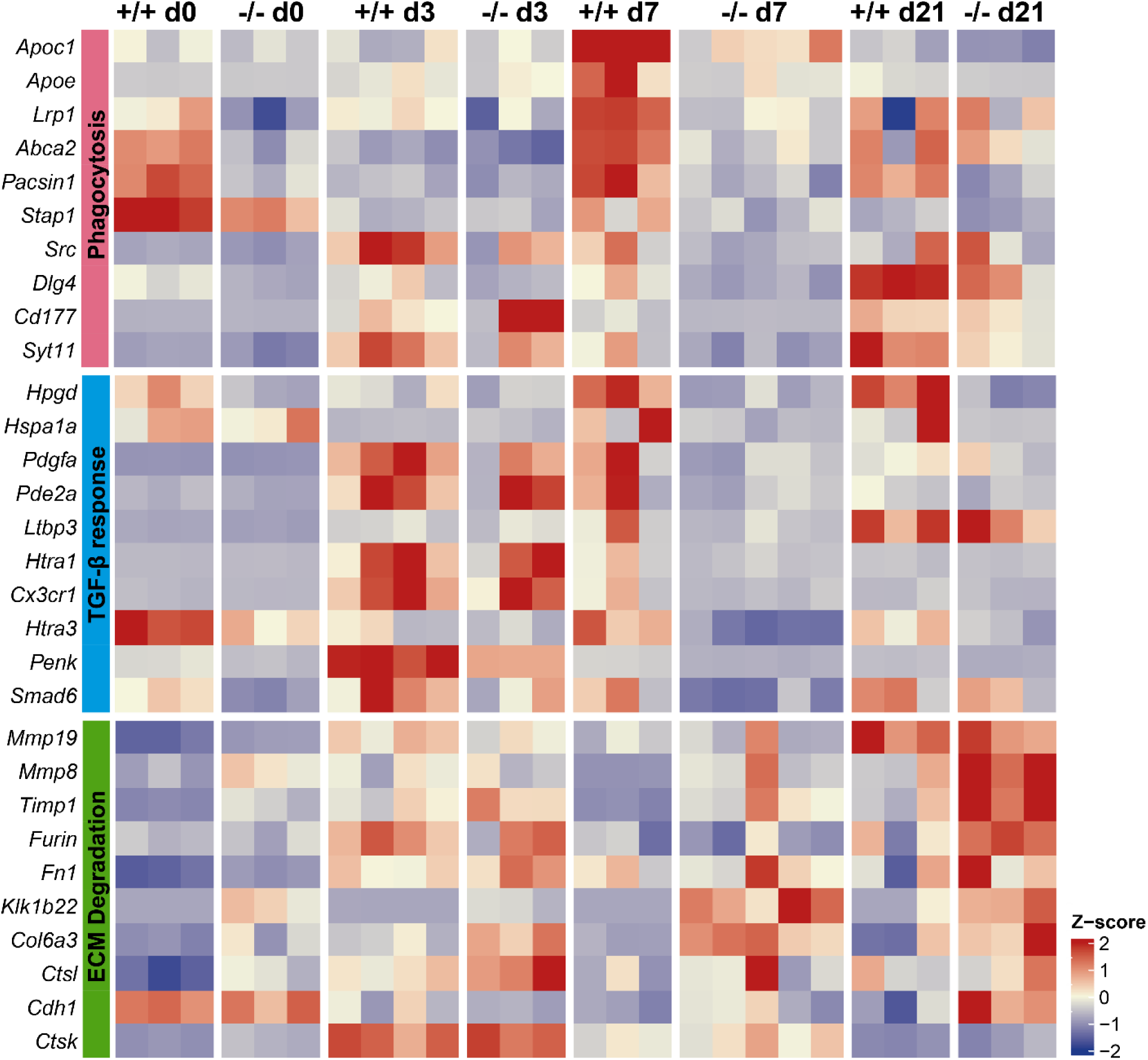
Absence of Siglec-F alters the transcriptional profile of RAMs. Gene set enrichment analysis (GSEA) was performed on bulk RNA-seq data from RAMs from *Siglecf^+/+^* and *Siglecf^-/-^* mice at days 0 (naïve), 3, 7, and 21 following PPE administration. The heatmap displays Z-score-normalized expression values for the top leading-edge genes from significantly enriched pathways (adjusted p-value < 0.05, normalized enrichment score > 1.5 or < -1.5).

On day 7, *Siglecf^-/-^* RAMs exhibited early transcriptional changes, including downregulation of TGF-β-responsive genes and phagocytosis-related genes. By day 21, the expression of extracellular matrix (ECM) degradation-related genes reached their highest levels, consistent with histological evidence of progressive tissue breakdown. These temporal gene expression patterns suggest that Siglec-F plays a role in controlling transcriptional programs related to tissue destruction, debris clearance, and repair in AMs, all of which are critical for maintaining lung structural integrity.

## Discussion

The studies reported here demonstrate that Siglec-F is crucial for suppressing the tissue-destructive effects of AMs. After PPE challenge, AMs were the predominant cell type associated with emphysema (Figure 1), and Siglec-F^+^ RAMs were the main AM subtype present (Figure 2). Loss of Siglec-F promoted airspace enlargement after PPE (Figure 3), and Siglec-F-deficient RAMs demonstrated significantly altered programming, consistent with more destructive phenotypes (Figure 4). Taken together, these studies identify important mechanistic links between Siglec-F expression and AM-mediated emphysematous alveolar destruction in mice.

We also observed decreased alveolar numbers in adult *Siglecf^-/-^*compared to *Siglecf^+/+^* mice in the absence of PPE challenge. This difference was not apparent on post-natal day 14, which marks the conclusion of peak alveologenesis in mouse post-natal lung development (19). Since Siglec-F deficiency did not appear to reduce the formation of alveoli, our findings suggest a potential role for Siglec-F and RAMs in sustaining alveolar maturation and integrity during later periods of lung growth. Although not examined at other pre-challenge timepoints here, potential roles for Siglec-F and RAMs could reflect programming differences that are evident after PPE-induced injury.

Chronic exposures to tobacco smoke, wood smoke, and other forms of air pollution are the leading causes of COPD worldwide. Emphysema develops over a course of decades in humans. Modeling COPD in mice is challenging, in part due to differences in their lung anatomy and lifespan (<2 years) compared to humans. Murine COPD models using cigarette smoke exposure require prolonged and repeated exposures that ultimately induce mild-to-moderate emphysematous phenotypes (20, 21). While these chronic models are valuable for translational relevance, their complexity and duration make it challenging to isolate and evaluate AM-specific mechanisms.

To address this, we used a single intratracheal challenge with PPE, which induces a robust emphysematous pathology. This acute model enables focused investigation of AM activation and damage-inducing responses in a relevant disease context. Additionally, we confirmed that PPE administration does not directly disrupt Siglec-F expression on AMs, allowing us to specifically assess the role of Siglec-F in modulating AM-mediated responses during lung injury. We anticipate that insights gained from this model will inform and enhance future studies using tobacco smoke exposure paradigms.

In our studies, we found that RAMs were the main AM subset present throughout the time course of injury. This predominance of RAMs contrasts with other injury models, such as bleomycin and LPS, where RecAMs outnumber RAMs during peak inflammation, with populations eventually resolving back to RAM dominance over time (8, 22). The relative absence of RecAMs in our model may reflect either the specific inflammatory context in our model or the fate of RecAMs after their initial recruitment. It is possible that in our model, severe epithelial injury impairs monocyte recruitment. Alternatively, RecAMs may undergo apoptosis or adopt RAM-like phenotypes (14, 23), rendering them indistinguishable by surface markers in flow cytometry by day 21.

Our data also identified a subset of neutrophils that express Siglec-F at day 3 post-injury, but their levels of Siglec-F expression are substantially lower than those of RAMs (Figure 2). Recent studies have identified Siglec-F expression on neutrophil subsets in various inflammatory contexts (24–28), including a report implicating Siglec-F⁺ neutrophils in promoting PPE-induced emphysema (29). While this suggests a pathogenic role for this transient population, our data show no difference in neutrophil numbers between Siglec-F-deficient mice and wild-type mice.

Instead, the most pronounced difference in leukocyte numbers between strains occurred in RAMs, which were markedly increased in Siglec-F-deficient mice at day 7. These findings raise the potential for a distinct and previously underappreciated role for Siglec-F in regulating RAM-driven tissue destruction.

Building on this evolving understanding of Siglec-F function in AMs, we previously showed that Siglec-F deficiency attenuated bleomycin-induced lung fibrosis (11). In contrast, our current findings reveal that Siglec-F deficiency exacerbates PPE-induced emphysematous lung injury, underscoring a protective role for Siglec-F in destructive lung disease. This apparent paradox reflects the distinct immunopathological mechanisms underlying fibrosis versus emphysema and supports our hypothesis that Siglec-F restrains AM activity in the latter’s context.

How Siglec-F exerts its immunoregulatory functions remains an active area of investigation. Like other CD33-related Siglec molecules, it contains an intracellular immunoreceptor tyrosine-based inhibitory motif (ITIM), typically associated with inhibitory signaling. In eosinophils, where Siglec-F function has been studied most extensively, its engagement induces apoptosis via a caspase-dependent mechanism that is independent of phosphorylation of ITIM (30, 31). Canonical ITIM-mediated suppression is well-characterized in human Siglecs such as Siglec-7 and Siglec-9, which are expressed on AMs (11, 32, 33). This mechanism does not fully apply to Siglec-F. In fact, a recent study using ITIM-deficient mutants has shown that Siglec-F signaling in eosinophils leads to cytokine production, and it occurs independently of ITIM, suggesting the involvement of alternative pathways (34). These findings underscore the context-dependent nature of Siglec signaling and point to broader roles in immune regulation and in AMs beyond classical inhibition.

Bulk RNA-seq analysis of sorted RAMs at key time points following PPE exposure provided mechanistic insights into how Siglec-F mediates this protective effect. Among the dysregulated pathways, ECM degradation emerged as a key axis. Notably, genes encoding for matrix metalloproteinases (MMPs) and their inhibitors, such as *Mmp19*, *Mmp8*, and *Timp1,* were all significantly increased in AMs from *Siglecf*^-/-^ compared to *Siglecf*^+/+^ mice (Figure 5). Increased expression of MMPs and their inhibitors is commonly observed in human COPD and in mouse models, consistent with what has been clinically described as a result of protease-antiprotease imbalance (35)

TGF-β signaling pathways were also significantly affected by *Siglecf* expression in AMs. TGF-β is a well-established regulator of the ECM in part through its suppression of MMP. Previous studies have shown that reduced TGF-β signaling leads to increased MMP expression and ECM degradation, contributing to emphysema development (36, 37). Impaired activation of TGF-β, as seen in mice lacking the β6 subunit of the αvβ6 integrin, leads to spontaneous alveolar enlargement, highlighting the critical role of TGF-β signaling in preserving alveolar integrity (38). Our transcriptomic data revealed downregulation of TGF-β signaling in Siglec-F-deficient RAMs, aligning with these prior findings.

Our RNAseq data also point to reduced expression of phagocytosis-related genes in Siglec-F-deficient RAMs, suggesting the potential for impaired apoptotic clearance as a contributing factor to sustained inflammation and tissue damage. AMs from COPD patients exhibit defective clearance of apoptotic cells and bacteria (39–42). In another PPE-induced murine emphysema model, Yoshida et al. showed that decreased efferocytosis worsened emphysema (43). AM efferocytosis mediates resolving lung inflammation after LPS challenge in a TGF-β-dependent (44). Given the corresponding downregulation of TGF-β and phagocytosis-related pathways in *Siglecf^-/-^* here, the potential role of Siglec-F as a checkpoint in RAMs to prevent tissue destruction in emphysema warrants further investigation.

In summary, our findings demonstrate that Siglec-F is more than just a marker - it actively regulates RAM-mediated homeostasis and effector functions. This aligns with previous reports highlighting Siglec family members as immunoinhibitory receptors that restrain inflammatory responses. In the context of emphysema, a disease driven by excessive protease activity and defective tissue repair, Siglec-F acts as a critical checkpoint, limiting RAM-mediated alveolar destruction. Specifically, by modulating genes involved in ECM degradation, phagocytosis, and tissue repair, Siglec-F suppresses deleterious RAM activities and preserves alveolar integrity. These insights enhance our understanding of immunoregulatory pathways controlled by Siglecs and highlight them as potential therapeutic targets in COPD, where current treatments fail to directly address the cellular drivers of alveolar damage.

## Supporting information

Supplemental materials_DEGs_list

Supplemental materials_GSEA_pathway_list

Supporting data values for all figures

## Author contributions

QY, WJJ, CME: conception, design, and writing, interpretation of data

QY, NH, YH, MO, BJS, TMB, RZB, ZL, BG, MK: acquisition, analysis, and interpretation of data

All authors reviewed the work critically, approved the final version, and agree to be accountable for all aspects of the work.

## Funding support

This research was supported by the following grants: NIH K99HL161446 (YH); NIH R01HL151630 (BJS); NIH F32HL172725 (MO); LongFonds, R01 HL141380 (MK); NIH R01HL14974, NIH R35HL140039 (WJJ); NIH R01HL130938 (CME, WJJ); NIH R01HL080396, VA I01BX005343, NIH P01HL162607 (CME).

Some data have been presented at the American Thoracic Society Conference in 2024 and at the American Association of Immunologists Conference in 2025.

This work is the result of NIH funding, in whole or in part, and is subject to the NIH Public Access Policy. Through acceptance of this federal funding, the NIH has been given a right to make the work publicly available in PubMed Central.

## Acknowledgement

We thank the Genomics Shared Resource (GSR) (RRID:SCR_021984) within the University of Colorado Cancer Center (P30CA046934) for their expert assistance in generating the RNAseq data. This work utilized the Alpine high performance computing (HPC) resource at the University of Colorado Boulder. Alpine is jointly funded by the University of Colorado Boulder, the University of Colorado Anschutz, and Colorado State University.

## Notes

**Conflict of interests**: The authors have declared that no conflict of interest exists.

### Competing Interest Statement

The authors have declared no competing interest.

