## Supplemental methods and figures for "Siglec-F Protects Against Elastase-induced Lung Inflammation and Emphysema in Mice"

### SUPPLEMENTAL MATERIALS

Qihua Ye[1], Naoko Hara[1], Yan Hu[2,3], Marika Orlov[1], Bradford J. Smith[4,5], Tonya M Brunetti[6], Rachel Z Blumhagen[7], Zhenghui Liu[1], Bifeng Gao[1], Melanie Königshoff[8,9], William J. Janssen[10,*], and Christopher M. Evans[1, *, †]

1. Division of Pulmonary, Allergy and Critical Care Medicine, University of Colorado Anschutz Medical Campus, Aurora, CO, United States.
2. Center for Perinatal Research, Nationwide Children's Hospital, Columbus, OH, United States.
3. Department of Pediatrics, The Ohio State University, Columbus, OH, United States.
4. Department of Bioengineering, University of Colorado Denver | Anschutz Medical Campus, Denver, CO, United States.
5. Division of Pediatric Pulmonary and Sleep Medicine, School of Medicine, University of Colorado Anschutz Medical Campus, Aurora, CO, United States
6. Department of Immunology and Microbiology, University of Colorado Anschutz Medical Campus, Aurora, CO, United States
7. Center for Genes, Environment and Health, National Jewish Health, Denver, CO, United States
8. Center for Lung Aging and Regeneration, Division of Pulmonary, Allergy, and Critical Care Medicine, Department of Medicine, University of Pittsburgh, Pittsburgh, PA, United States
9. Geriatric Research Education and Clinical Center at the VA Pittsburgh Healthcare System, Pittsburgh, PA, United States
10. Department of Medicine, National Jewish Health, Denver, CO, United States

* These authors contributed equally to this publication

† Corresponding author:

Christopher M. Evans, PhD

303-724-9287

#### Supplemental Methods

##### Mice

Mice were purchased from the Jackson Laboratories (Bar Harbor, ME). Housing rooms are maintained at 22°C, 30-40% humidity, and a 14/10 (h/h) light-dark cycle. *Siglecf*$^{-/-}$ mice on a congenic C57BL/6J background and maintained by breeding with C57BL/6J. *Siglecf*$^{+/+}$ littermates and purchased C57BL/6J mice were used. A mix of male and female mice aged 12-18 weeks at dosing was used.

##### Induction of lung inflammation and emphysema

Mice were anesthetized with isoflurane and instilled with porcine pancreatic elastase (PPE, Sigma, cat. E7885, lot # 1003571181) oropharyngeally (20 U/kg).

##### Lung function measurement

Lung function was measured using a flexiVent FX1 (Scireq, Montreal, Quebec, Canada). Mice were anesthetized with urethane (2.0 g/kg, i.p.), tracheostomized with a blunt beveled 18 Ga Luer stub, and ventilated with a respiratory rate of 150 breaths/min and a tidal volume of 10 ml/kg against 3 cm $H_2O$ positive end expiratory pressure. Lung function was measured by first performing a 6-s "deep inflation" to 30 cm$H_2O$. Then, we performed a 1.25-s single-frequency forced oscillation signal at 150 breaths/min (2.5 Hz) and fit the single-compartment model to determine respiratory system resistance (Rrs) and respiratory system elastance (Ers). A 3-s broadband low-frequency forced oscillation signal was used to measure input impedance from 1-20.5 Hz and to calculate constant phase model parameters, including pulmonary system elastance (*H*), tissue damping (*G*), and Newtonian (airways) resistance (*Rn*). Last, we performed a 16-s stepwise pressure-volume (PV) loop with an onset pressure of 3 cm$H_2O$ and a maximum pressure of 30 cm$H_2O$ to calculate quasi-static compliance (Cst, the slope of the quasi-static points at 5 cm$H_2O$ on the expiratory limb). Inspiratory capacity (IC) measurement was derived from the volume at 30 $cmH_2O$, the highest point of the PV loop. This sequence was performed three times, and the average was taken to report for each mouse.

**Histology**

After puncturing the diaphragm and performing a bilaterial thoracotomy, lungs were inflated and fixed with 1% low-melting agarose and 4% PFA at a pressure of 25 cm $H_2O$ through the tracheal cannula. After 72 h of immersion fixation in 4% PFA, extrapulmonary tissues were removed, and the total lung volume was determined using Archimedes principle. Subsequently, the whole lung was embedded in 3% agar with random lobe orientations. The lung-containing agar was cut into 2 mm slabs, and the even or odd slabs were randomly selected for paraffin embedding. Two sequential 5 µm H&E-stained serial sections were obtained from the paraffin-embedded cassettes for stereology. Images were taken on Olympus VS120 Scanner at 20X.

**Stereological analysis**

Stereological analysis included 1) quantification of alveolar number and 2) airspace enlargement. Analyses were performed using VIS (2017.7.3, Visiopharm, Hørsholm, Denmark). The physical dissector was used to estimate alveolar number. Two serial whole-slide images were superimposed, and systematic uniform random sampling (SURS) was conducted at 40X digital magnification with an area sampling fraction was 10% for day 0 and 20% for day 21. The randomly sampled fields of view were displayed side-by-side with superimposed 150 µm x 150 µm counting frames. Alveoli were counted when the mouth of the alveoli was present in one counting frame but not the other, and the alveolar mouth did not touch the exclusion lines (Fig. S3A). The final alveolar count per mouse was calculated using established methods (1), with tissue shrinkage correction applied (2).

$$\text{Total Alveolar Number} = \left( \frac{\#\ \text{septal breaks}}{2 \cdot (\#\ \text{sections} \cdot V_d)} \right) \cdot V_l$$

$V_d$ : Dissector volume, 150 * 150 * 5 μm = 112,500 $\mu m^3$.

$V_l$: Total lung volume, measured using Archimedes principle.

To quantify airspace enlargement, chord lengths were measured. The SURS fraction at 20X was 20% for days 0, 1 and 3, and 40% for days 7 and 21. The reference line was 150 μm (Fig S3B, blue) and the remaining length of the counting line is a guard zone, which is 300 μm (red). Working left to right, the start of the chord was marked at the septal-air interface and the airspace chord was terminated at the next septa. Chords were terminated, but not started, in the guard zone. The final reported mean chord lengths were the average of all individual chord lengths.

**Murine macrophage isolation**

At 0, 1, 3, 7, and 21 days after PPE instillation, lung lavage was performed, and tissues were collected. Mice were euthanized by exsanguination. With the left lung clamped at the mainstem bronchus, the right lung was lavaged by instilling and removing 0.5 ml three times with PBS containing 0.5 mM EDTA (1.5 ml total). Lung lavage was used for hemocytometer counts, Giemsa stain differentials, flow cytometry, and FACS.

For RNAseq studies, lung lavage was performed as above, except that 1 ml PBS/EDTA was used to lavage whole lungs three times (3 ml total). Lung lavage was centrifuged at 300 x g for 8 min at 4°C, and cells were used for FACS and RNA isolation.

**Murine macrophage flow cytometry and sorting**

Lung lavage was collected as above and labeled with the following markers using an antibody dilution of 1:200 (v/v) in the presence of TruStain FcX™ anti-CD16/32 (Clone 93, Biolegend, San Diego, CA, cat. 101320): CD45 FITC (Clone 30-F11, Biolegend, cat. 103108), CD64 PE-Cy7 (Clone X54-5/7.1, Biolegend, cat. 139314), CD88 APC (Clone 20/70, Biolegend, cat. 135808), Ly6G Pacific Blue and BV605 (Clone 1A8, Biolegend, cat. 127612/127639), CD11b APC-Cy7 (Clone M1/70, Biolegend, cat. 101226), CD11c PerCP (Clone N418, Biolegend, cat. 117326), and Siglec-F PE (Clone S17007L, Biolegend, cat. 155506) at 4°C for 45 min. The cells were centrifuged at 300 x g for 8 min, washed in FACS buffer, and re-suspended in 500 µl FACS buffer (Biolegend, cat. 420201). Before sorting, one drop of NucBlue (Invitrogen, Waltham, MA, cat. R37606) or DAPI (Biolegend, cat. 422801) was added to the cells to exclude dead cells. AMs were gated as $CD45^{+}CD64^{+}CD88^{+}Ly6G^{-}$ cells, and recruited and resident AMs were separated based on CD11b and CD11c expression, respectively. FACS was performed on a Sony Biotechnology SY3200 sorter. Flow cytometry was performed on a BD LSR Fortessa analyzer.

**Bulk RNA-seq and bioinformatics**

RNA was isolated from sorted cells using micro RNA kit (Qiagen, Germantown, MD). RNA purity, quantity, and integrity were determined by NanoDrop (ThermoFisher Scientific, Waltham, MA) and TapeStation 4200 (Agilent, Santa Clara, CA) analyses prior to RNA-seq library preparation. MGIEasy RNA Library Prep Set (3.1, cat. 1000006384) was used to generate libraries with an input of 100 ng of total RNA. Paired-end sequencing reads (100 bp) were generated on the DNBSeq-T7 sequencer (MGI/Complete Genomics, San Jose, CA), targeting approximately 80 million total reads per sample. Raw fastq files were trimmed using cutadapt (4.2) and aligned to the reference genome GRCm38/mm10 using STAR (2.7.10b). Picardtool (2.27.5) was used to collect strandedness information for each sample. Raw gene-level counts were filtered to remove genes with low expression, retaining only those with >10 raw counts in at least 10 of the 27 total samples. This strategy removes genes with limited and inconsistent expression across the dataset while preserving those robustly detected across multiple samples. Normalized counts were acquired using DESeq2 (1.38.2) in R (4.2.2), followed by shrinkage of effect size. Differential expression analysis was performed using DESeq2 (1.38.2) in R (4.2.2). Gene set enrichment analysis (GSEA) was performed using fgsea (1.24.0). The genes were ranked and collectively mapped to the following gene sets: H (Hallmark), C2CP (Curated gene set canonical pathways), and C5BP (Gene ontology biological processes). Featured pathways were visualized using ComplexHeatmap (3.22) in R (4.2.2).

**Statistical analysis**

For non-RNAseq data, statistical analyses were conducted using Prism 10.2.3 (GraphPad, La Jolla, CA). Two-sample comparisons were performed with unpaired two-tailed *t*-tests or Mann-Whitney U-tests where appropriate based on the distribution. For multiple comparisons, one-way ANOVA with a Dunnett post-hoc correction or a Kruskal-Wallis test with Dunn’s post-hoc correction was used.

For RNA-seq data, differentially expressed genes (DEGs) were identified using a threshold of adjusted p-value of $<0.05$ and $\log_2$FC of $> 1$ or $< -1$. Significant pathways identified through GSEA were defined by an adjusted p-value $< 0.05$ and normalized enrichment scores (NES) $>$ 1.5 or $< -1.5$.

**Supplemental figures and legends**

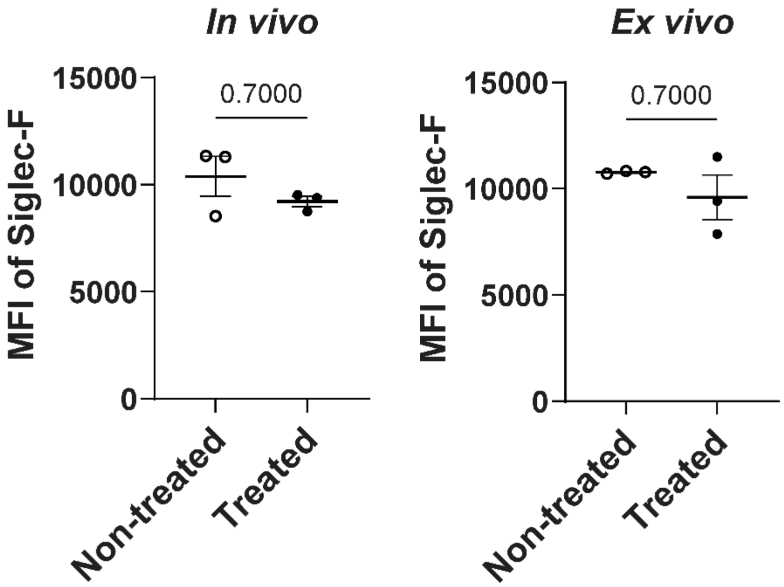


**Figure E1. Siglec-F surface expression remains unchanged after exposure to PPE in vivo and in vitro.** Siglec-F surface expression following 1 h incubation with PPE in vivo and in vitro. Statistical analysis was performed using Mann-Whitney test. Data are presented as mean ± SEM.

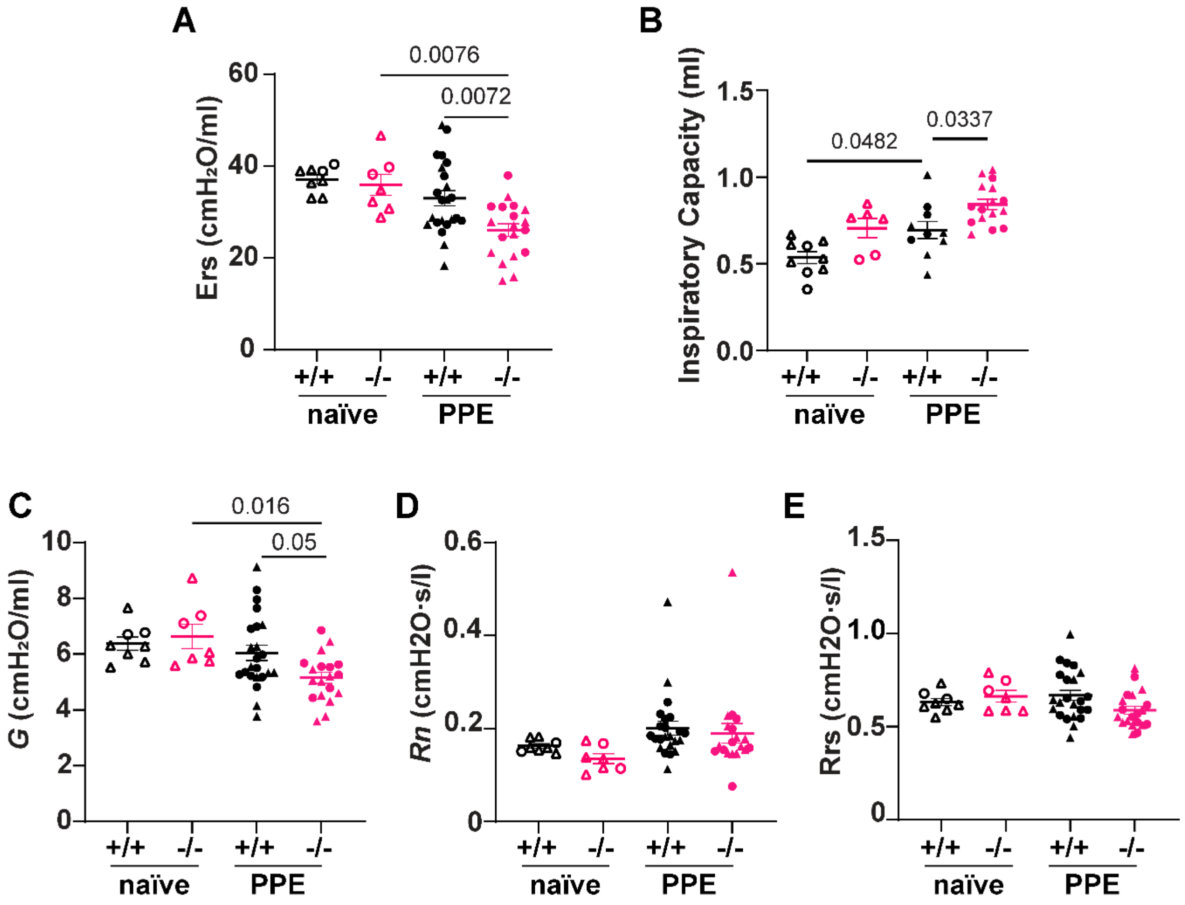


**Figure E2. Lung mechanics parameters. (A)** respiratory system elastance (Ers), **(B)** inspiratory capacity **(C)** tissue damping (*G*), **(D)** Newtonian (airways) resistance (*Rn*), **(E)** Respiratory system resistance (Rrs). Triangles denote female mice; circles denote male mice. Data are shown as mean ± SEM. One-way ANOVA with Dunnett's post hoc test was performed.

A

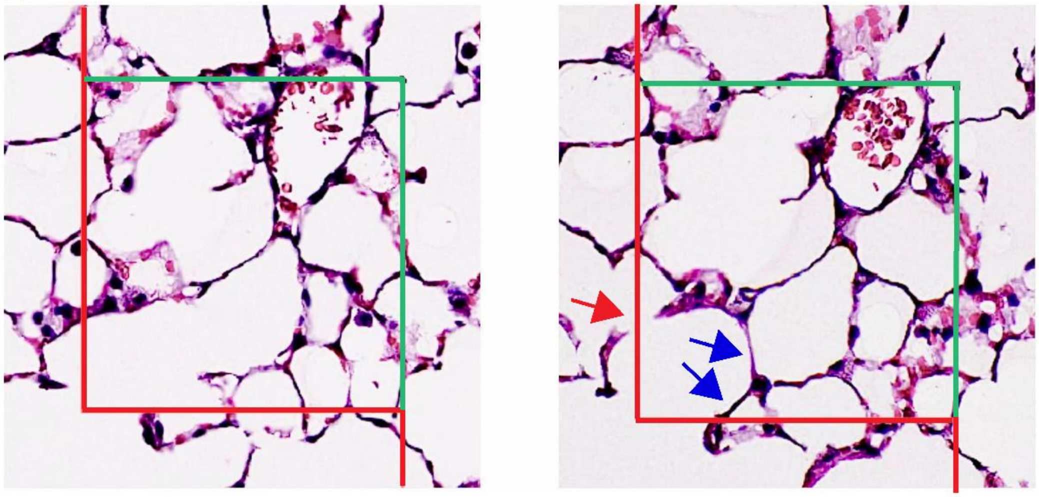

B

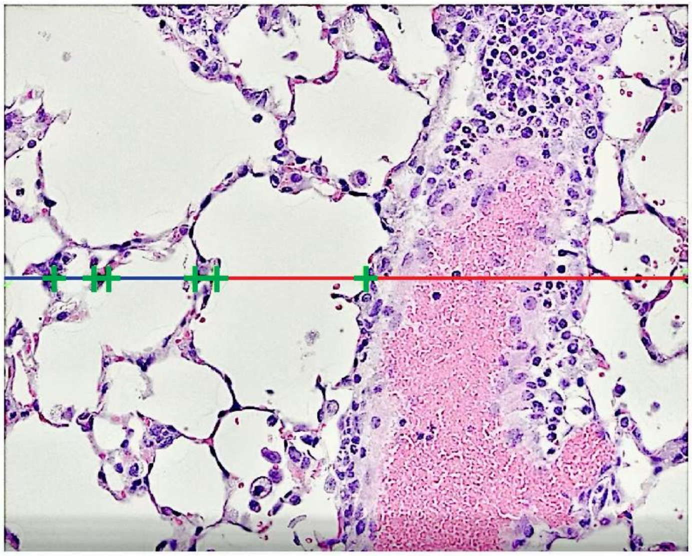

**Figure E3. Identification of septal breaks for alveolar quantification and intercepts for chord length measurement. (A)** Blue arrows indicate septal breaks identified across serial sections. The red arrow marks a septal break excluded from counting due to its intersection with the red line. **(B)** Green markers indicate points where alveolar septa intersect the counting line.

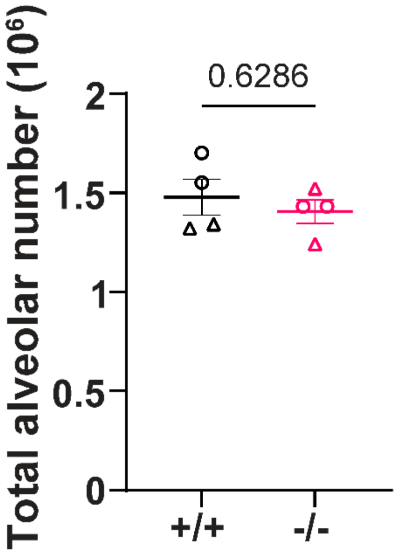


**Figure E4. Siglec-F deficiency does not lead to decreased alveoli post-natal day 14 (p14).** Quantification of alveoli in *Siglecf*$^{+/+}$ and *Siglecf*$^{-/-}$ mice on p14. Data are shown as mean ± SEM. Mann-Whitney test was performed.

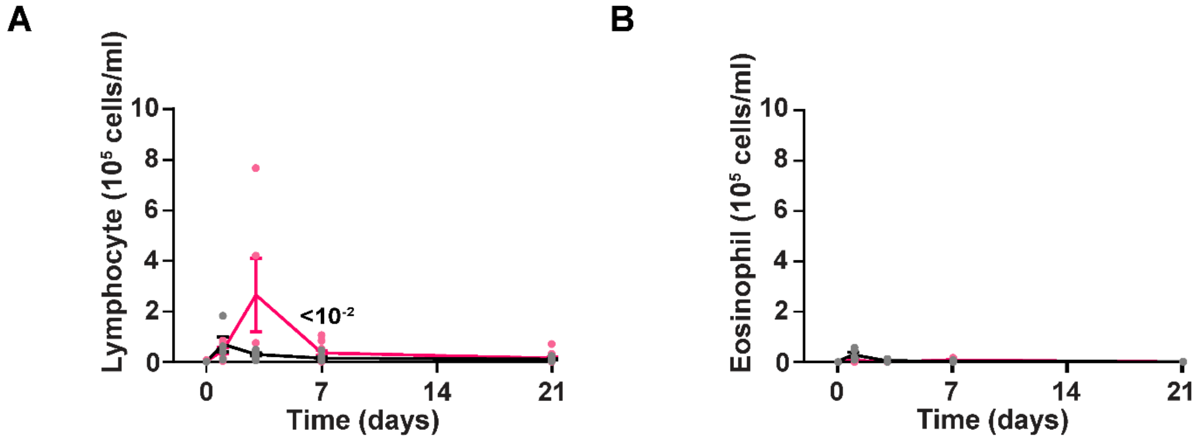


**Figure E5. Siglec-F deficiency leads to transient increase in lymphocytes and no difference in eosinophils.** Quantification of **(A)** lymphocyte and **(B)** eosinophil in lung lavage from *Siglecf*$^{+/+}$ (black) and *Siglecf*$^{-/-}$ mice (magenta) (n = 4-10 per time point). Data are shown as mean ± SEM. Statistical analyses between *Siglecf*$^{+/+}$ and *Siglecf*$^{-/-}$ mice at each time point were performed using Mann-Whitney test.
